# Ectomycorrhizal fungi induce systemic resistance against insects on a non-mycorrhizal plant in a CERK1-dependent manner

**DOI:** 10.1101/852640

**Authors:** Kishore Vishwanathan, Krzysztof Zienkiewicz, Yang Liu, Dennis Janz, Ivo Feussner, Andrea Polle, Cara H. Haney

## Abstract

Below-ground microbes can induce systemic resistance (ISR) against foliar pests and pathogens on diverse plant hosts. The prevalence of ISR among plant-microbe-pest systems raises the question of host specificity in microbial induction of ISR. To test whether ISR is limited by plant host range, we tested the ISR-inducing ectomycorrhizal (ECM) fungus *Laccaria bicolor* on the non-mycorrhizal plant *Arabidopsis*. We found that root inoculation with *L. bicolor* triggered ISR against the insect herbivore *Trichoplusia ni* and induced systemic susceptibility (ISS) against the bacterial pathogen *Pseudomonas syringae* pv. tomato DC3000 (*Pto*). We found that *L. bicolor-*triggered ISR against *T. ni* was dependent on jasmonic acid (JA) signaling and salicylic acid (SA) biosynthesis and signaling. We found that heat killed *L. bicolor* and chitin are sufficient to trigger ISR against *T. ni* and ISS against *Pto* and that the chitin receptor CERK1 is necessary for *L. bicolor-*mediated effects on systemic immunity. Collectively our findings suggest that some ISR responses might not require intimate co-evolution of host and microbe, but rather might be the result of root perception of conserved microbial signals.

## INTRODUCTION

Plants associate with complex communities of microorganisms. Interplay of host and microbial genotype determine whether the outcome of specific plant-microbe interactions is beneficial or detrimental to plant health^1–3^. Plants possess receptors to sense potential pathogens including transmembrane pattern-recognition receptors that recognize conserved microbe-associated molecular patterns (MAMPs), and intracellular receptors that directly or indirectly recognize effectors to help limit pathogen growth^4,5^. MAMP perception by plant receptors triggers pattern-triggered immunity (PTI) responses including a reactive oxygen species (ROS) burst, Mitogen-Activated Protein Kinase (MAPK) signaling, ion influx and callose deposition^6–9^. In addition to local immune mechanisms, root-associated microbes can induce systemic resistance (ISR) against a diverse spectrum of above-ground threats^2,10^. While the mechanisms by which plants perceive MAMPs and effectors are well understood, there is a more limited understanding of the mechanisms by which plant perceive the microbes that trigger ISR.

In response to perception of microbes or pathogens, plants activate systemic defense mechanisms including commensal-triggered ISR and pathogen-triggered systemic acquired resistance (SAR)^11,12^. While both SAR and ISR confer resistance to pests and pathogens, they promote resistance through distinct mechanisms^2^. SAR occurs upon local perception of a pathogen, MAMP, or effector and results in systemic induction of SA-dependent gene expression and accumulation of secondary metabolites^13,14^. Unlike SAR, ISR is associated with small or no changes in systemic gene expression and phytohormone levels^15,16^. Additionally, while SAR is dependent on the SA signaling protein NPR1^13^, the pathways required for ISR can vary between systems^17,18^.

ISR triggered by a single microbial strain can be effective across diverse hosts and against diverse pathogens. For instance, *Pseudomonas simiae* WCS417 was isolated from barley fields^19^ and can trigger ISR on carnation, tomato, wheat and *Arabidopsis*^20–22^. Similarly, *P. defensor* WCS374 and *P. capeferrum* WCS358, which were isolated from potato fields, trigger ISR in radish and *Arabidopsis* respectively^21,23^. That ISR by a single bacterial strain is effective against phylogenetically diverse plants suggests that ISR is likely dependent on broadly conserved plant perception and signaling mechanisms.

In addition to interactions with rhizobacteria, most vascular plants (>90%) form beneficial symbioses with arbuscular mycorrhizal (AM) fungi or ectomycorrhizal (ECM) fungi^24^. Both AM and ECM fungi can induce ISR on their host plants against foliar pests and pathogens^25–29^. AM symbiosis requires the Common Symbiotic Pathway (CSP) for initiation of successful symbiosis^30^; recent evidence suggests that ECM symbiosis may also depend on the CSP^31^. It is unknown whether mycorrhizal-triggered ISR is dependent on the CSP and is dependent on the same or distinct mechanisms from rhizobacterial-triggered ISR.

Here, we tested the hypothesis that ISR occurs independently of mutualistic symbiosis or host specificity. To address our hypothesis, we tested whether *Laccaria bicolor*, an ECM fungus that colonizes a wide range of tree host species^32^ could trigger ISR on *Arabidopsis thaliana*, a non-mycorrhizal plant^24^. *Arabidopsis* lacks the CSP and so allows for direct testing of whether ISR by mycorrhizal fungi is dependent on the CSP. We rationalized that if ISR is the result of a mutualistic symbiosis, then we should not observe ISR by *L. bicolor* on *Arabidopsis*. In contrast, if ISR is the result of plant perception of a general microbial factor, we might observe ISR by *L. bicolor* on *Arabidopsis*.

Consistent with the hypothesis that ISR against herbivores can occur independently of highly co-evolved mutualism, we found that *L. bicolor* can trigger ISR on *Arabidopsis* against caterpillars of the generalist herbivore *Trichoplusia ni* and induce systemic susceptibility (ISS) against *P. syringae* pv. tomato DC3000 (*Pto*). We found that *L. bicolor*-triggered ISR against *T. ni* and ISS against *Pto* requires CERK1-dependent chitin perception. We show that chitin and heat-killed fungi are sufficient to modulate systemic immunity and this response is distinct from root perception of bacterial MAMPs. Collectively this work shows that chitin perception by roots is sufficient to modulate systemic plant immunity and that ISR against herbivores might be a general response to perception of chitin, independent of host adaptation.

## RESULTS

### *L. bicolor* triggers ISR on *Arabidopsis* against *T. ni*

To test the hypothesis that ISR occurs independently of mutualist symbiosis, we treated the non-mycorrhizal plant *Arabidopsis thaliana* with the ECM fungus *Laccaria bicolor* and measured ISR against the generalist herbivore *Trichoplusia ni*^17,33^. The roots of 9-day-old *Arabidopsis* seedlings were treated with *L. bicolor* (Methods), and the rosettes were challenged with *T. ni* larvae 3 weeks later. After 1 week of feeding, we found that *T. ni* larvae that fed on *L. bicolor*-treated *Arabidopsis* had a 27% reduction in larval weight compared to buffer-treated plants (Figure 1a, *p* < 0.006). The observed *L. bicolor*-induced ISR is similar in magnitude to previously described bacterial treatments that induce ISR against *T. ni* on *Arabidopsis*^17,34,35^.

**Figure 1.**
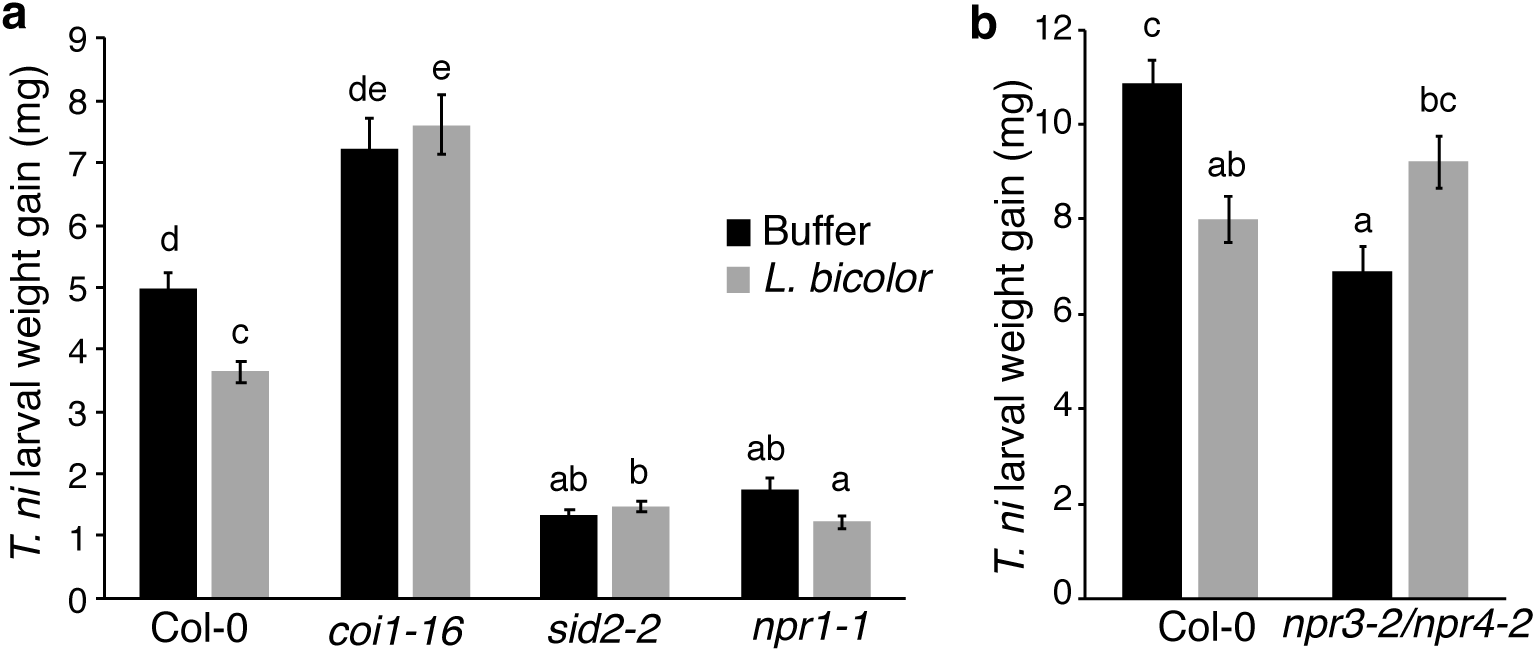
*L. bicolor* treatment of *Arabidopsis* roots induces systemic resistance against *T. ni* via JA signalling and negative regulators of SA signalling. **a**, ISR triggered by *L. bicolor* is dependent on the JA-Ile receptor *COI1* and SA biosynthesis via *SID2*. For *coi1-16* and *npr1-1*, n = 4 experiments and for *sid2-2*, n = 3 experiments with at least 20 caterpillars per treatment. **b**, ISR triggered by *L. bicolor* is dependent on *NPR3/4*. n = 3 experiments with at least 20 caterpillars per treatment. Statistical significance between treatments was determined by performing two-way ANOVA of the linear mixed model with Tukey’s test. Different letters indicate significance at *p* < 0.05 and the error bars indicate SE.

These data suggest that ISR by the ECM fungus *L. bicolor* does not require host colonization. As ISR can occur by distinct mechanisms against different herbivores and pathogens, we sought to compare and contrast *L. bicolor* induced ISR to other well-studied mechanisms of ISR on *Arabidopsis*.

### ISR by *L. bicolor* requires SA and JA signalling

ISR against herbivores triggered by beneficial *Pseudomonas* spp. is dependent on JA signalling^17,34,35^. To determine if *L. bicolor*-induced ISR against *T. ni* is also dependent on JA signalling, we tested ISR by *L. bicolor* on the JA-Ile receptor mutant, *coi1-16*. As previously shown, we found that the *coi1-16* mutant has enhanced susceptibility to *T. ni*^17,36^. We found that the *coi1-16* mutant did not show reduced caterpillar weight gain after *L. bicolor* treatment (Figure 1a). Therefore, like ISR by *Pseudomonas* spp. against herbivores, ISR by *L. bicolor* in *Arabidopsis* is dependent on JA-Ile perception via *COI1*.

ISR against herbivores triggered by *Pseudomonas* spp. is dependent on SA biosynthesis via *SID2* and is independent of SA signalling via *NPR1*^17^. We found that a *sid2-2* mutant (deficient in SA biosynthesis) did not show ISR in response to *L. bicolor* (Figure 1a). In contrast, *L. bicolor* treatment of the SA signalling mutant *npr1-1* resulted in a 30% reduction in *T. ni* weight gain (Figure 1a, *p* < 0.11). Consistent with previous studies, we found that the *sid2-2* and *npr1-1* mutants show enhanced resistance to *T. ni*^17,33^. Though not statistically significant, the ISR induced on the *npr1-1* mutant was similar in magnitude to wild-type plants (Figure 1a). *T. ni* larvae feeding on an *npr1-1* mutant are extremely small leading to challenges in quantifying further reduction in weight^17,33^. Given the consistent and similar magnitude in resistance by *L. bicolor* treatment with *npr1-1* mutant, it seems likely that *L. bicolor* triggered ISR is independent of or partially dependent on *NPR1*. Overall, these data show that similar to ISR against *T. ni* by *P. simiae* WCS417, ISR by *L. bicolor* requires both JA signalling and SA biosynthesis^17^.

We found that ISR by *L. bicolor* against *T. ni* is dependent on *SID2* but partially dependent or independent of the positive regulator, *NPR1*. This raises the possibility that negative regulators of SA signalling are required for ISR by *L. bicolor*. Ding et al. (2018) reported that *NPR3* and *NPR4* act as SA receptors that repress SA defence gene expression in an *NPR1* independent manner. To determine whether *L. bicolor* mediated ISR is dependent on SA transcriptional co-repressors, the *npr3-2 npr4-2* (*npr3/4*) double mutant, was tested for ISR. We found that in contrast to wild type plants, *L. bicolor* treatment of an *npr3/4* mutant resulted in a significant increase in *T. ni* larval weight gain (Figure 1b, *p* < 0.003). The variance in the weight gain of *T. ni* larvae feeding on control plants relative to the data in Figure 1a for this and subsequent experiments can be attributed to spatial and temporal factors^17,36^; replicates of the same experiment were done in the same location and over a relatively short period of time while distinct experiments were done over longer time scales and in multiple locations leading to variance in the absolute weight gain of the controls^17,36^ (Supplementary Figure 1). These data indicate that *L. bicolor*-triggered protection of *Arabidopsis* from caterpillar feeding is dependent on the negative regulators of SA signalling, *NPR3/4*.

### *L. bicolor* induces systemic defence-gene expression

Previous reports of ISR by beneficial *Pseudomonas* against insect herbivores are associated with a low-level priming of expression (1.5-to 3.5-fold) of JA-Isoleucine (JA-Ile) dependent genes^17,38^. In contrast, SAR is associated with strong priming (4-to 10-fold) of SA-dependent gene expression in distal leaves^39,40^. To determine if ISR by *L. bicolor* is associated with priming of systemic defence-gene expression, we performed RNAseq to analyse gene expression in *Arabidopsis* leaves after root treatment with *L. bicolor* (Figure 2a and Supplementary Table 1). In response to *Arabidopsis* root treatment with *L. bicolor*, we found a significant increase in systemic expression of hormone-responsive genes including JA- and Ethylene-dependent genes (Figure 2a). We also found an increase in expression of genes involved in secondary metabolism and chitin responses (Figure 2a). We found that *L. bicolor* treatment of roots resulted in significant induction in shoots of JA-Ile marker genes including *VSP1, VSP2, MYC* and MAMP-responsive genes including *MYB51* and *WRKY70* (SA), which have been shown to be induced by root treatment with *Pseudomonas* spp.^17,38^ (Figure 2b). The initial RNAseq experiments were performed on gnotobiotic plants growing on solid media (Methods); to determine if the responses we saw were further enhanced after challenged with herbivores, we grew plants in soil and treated with *L. bicolor.* In soil-grown plants, we found no significant systemic changes in gene expression in response to *L. bicolor* treatment by RNAseq (Methods) or qPCR (Figure 2c). While challenging leaves with *T. ni* resulted in significant induction of JA-Ile-dependent defence-gene expression relative to unchallenged plants, we did not find any further enhancement of gene expression up *L. bicolor* treatment (Figure 2c). The lack of significant changes in soil grown plants during ISR is consistent with previous reports of no or slight priming of systemic gene expression^17,22^. The slight increase in systemic gene expression after *L. bicolor* treatment is consistent with an ISR-like rather than with a SAR-like mechanism (Figure 2).

**Figure 2.**
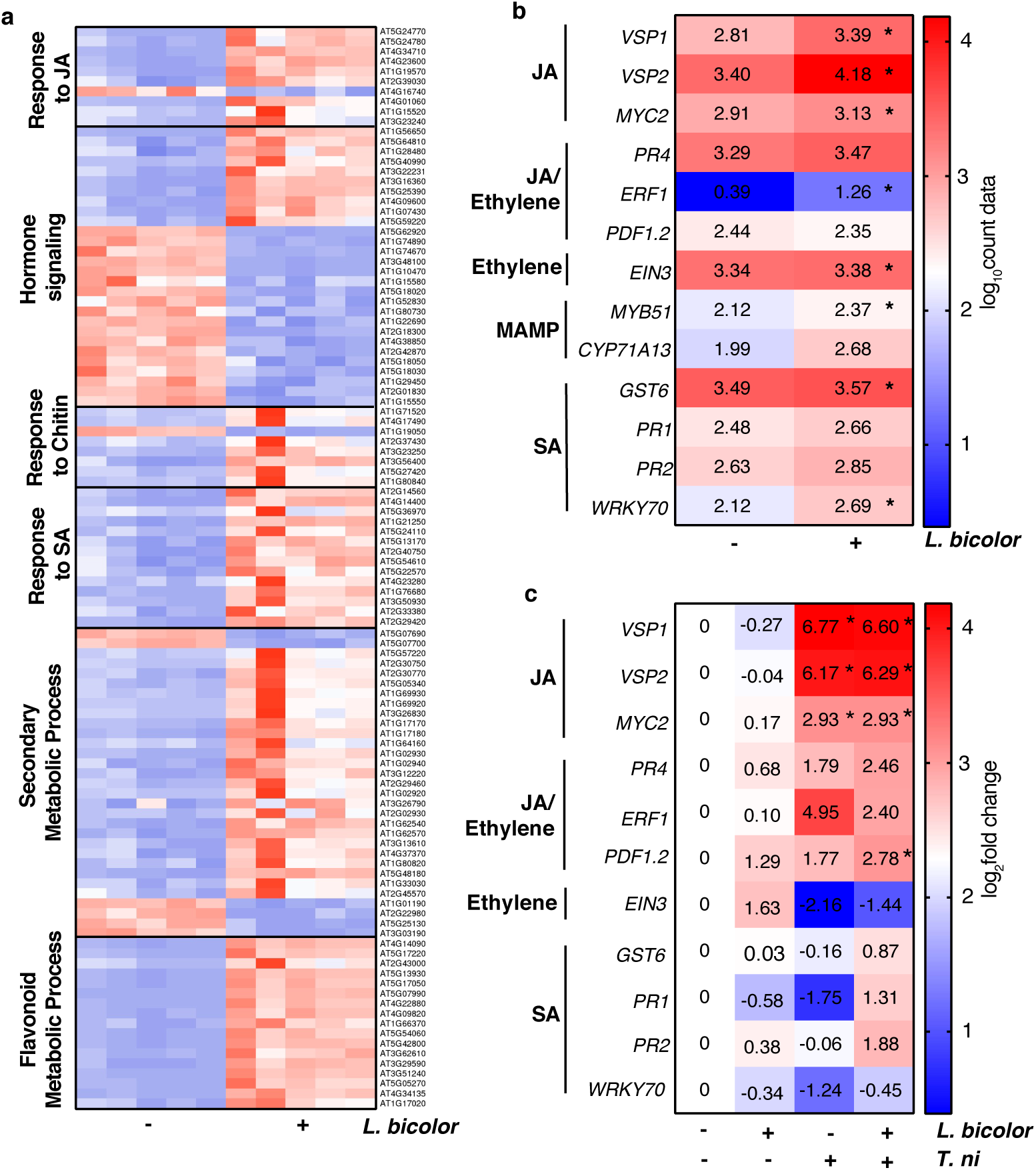
*L. bicolor* induces systemic hormone and defence-gene expression in *Arabidopsis*. **a**, Heat map showing row Z-score of highly upregulated and downregulated genes from specific gene ontologies in Col-0 leaves after root treatment with buffer or *L. bicolor.* Data from 5 independent replicates are shown. **b**, RNAseq data showing *Arabidopsis* hormone- and defence-regulated genes in response to *L. bicolor*. Data show log_10_ (count data) of defence genes in samples collected from 5 independent experiments (n = 5). Adjusted *p* value was calculated using Wald test. Asterisks denote statistical significance at *p* < 0.05. **c**, qPCR showing expression of defence-related genes in leaves after *L. bicolor* treatment of roots. CT values of defence genes obtained using qRT-PCR were normalized to the house keeping gene *EIF4A*. Data show log_2_ relative fold change [log_2_ (2^-(ΔΔCT)^)] in samples collected from six independent experiments (n = 6) normalized to the mock-inoculated plants. Statistical analysis was performed for the [log_2_ (2^-(ΔΔCT)^)] values using two-way ANOVA with Tukey’s HSD test.

SAR is associated with elevated SA levels in distal tissues^41^, while ISR is not associated with significant changes in systemic hormone accumulation^2^. Phyto-hormone analyses was performed using leaves from the same soil grown plants used for the qRT-PCR experiment above. We found that similar to the qRT-PCR results, *L. bicolor* treatment of *Arabidopsis* roots resulted in no significant increase in accumulation of the phytohormones SA, ABA and JA-Ile (Supplementary Figure 2). As expected, plants challenged with *T. ni* had a significant increase in JA levels; however, these effects were not further enhanced after *L. bicolor* treatment. Collectively, these data indicate that *L. bicolor* triggers systemic resistance against herbivory with moderate priming of defence gene expression and no significant accumulation of phytohormones in *Arabidopsis* leaves.

Low-molecular weight secondary metabolites contribute to plant defence and to systemic resistance against invading pathogens and other threats^29,35,42^. Our RNAseq data showed that *L. bicolor* treatment resulted in significant upregulation of the camalexin biosynthesis genes *PAD3* and *CYP71A13* (Figure 2 and Supplementary Table 1). To determine whether secondary metabolites accumulate in systemic leaves after *L. bicolor* treatment, targeted metabolite analysis was performed with *Arabidopsis* plants with and without *L. bicolor* treatment and *T. ni* feeding. We found that similar to *Arabidopsis* treatment with beneficial *Pseudomonas* spp.^35^, camalexin levels in leaves were significantly higher in *L. bicolor*-treated plants than in buffer-treated plants (Figure 3a, *p* < 0.05). Camalexin levels were increased by *T. ni* challenge, and *L. bicolor*-pretreated plants accumulated higher levels of camalexin after *T. ni* challenge than the buffer-treated controls (Figure 3a). In contrast, we did not observe significant accumulation of indolic glucosinolates like glucobrassicin and their by-product raphanusamic acid, which are involved in pathogen- and insect-triggered defense^43,44^ (Supplementary Figure 3). Camalexin is derived from the tryptophan pathway, which is also the precursor for many metabolites with insecticidal properties^42^. The mutant *cyp79b2/b3*, which is impaired in the biosynthesis of indole-3-acetaldoxime (IAOx) and camalexin from tryptophan, was tested for *L. bicolor* induced resistance against *T. ni* larvae. Pre-treatment of the *cyp79b2/b3* mutant with *L. bicolor* did not result in significant decrease in *T. ni* larval weight (Figure 3b). This indicates that the IAOx pathway is required for *L. bicolor* triggered systemic protection against herbivore damage.

**Figure 3.**
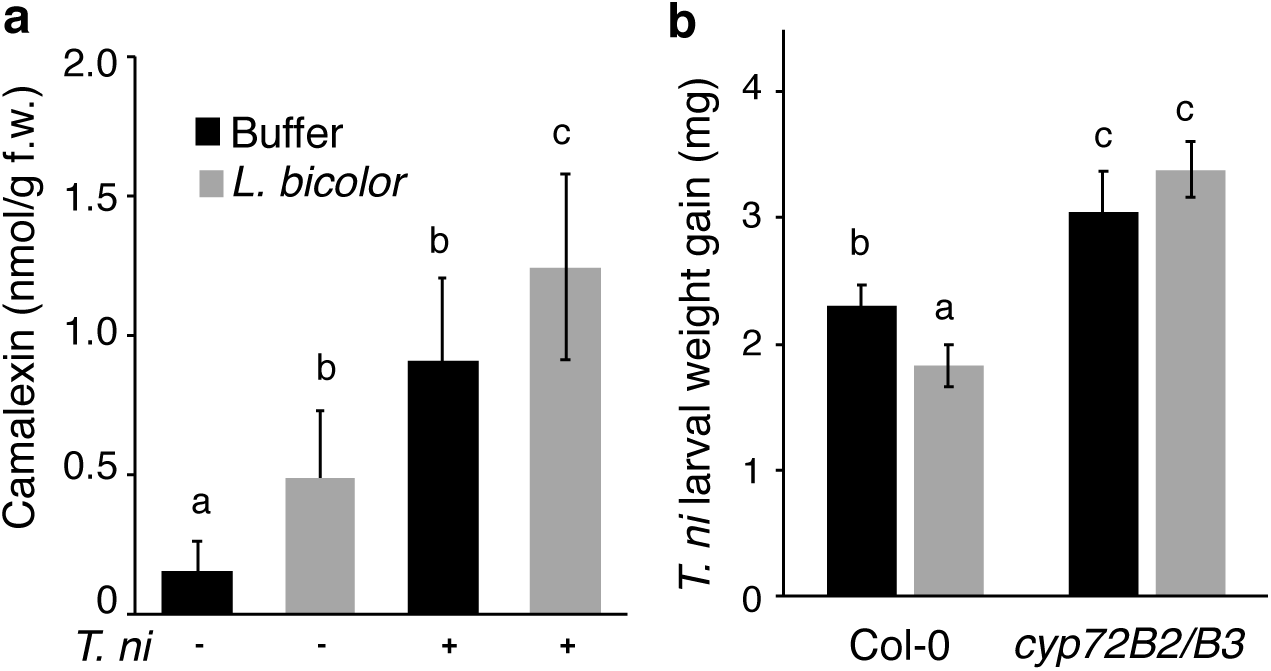
*L. bicolor* treatment of *Arabidopsis* roots results in increased camalexin in shoots. **a**, Leaves were harvested from *Arabidopsis* plants treated with buffer or *L. bicolor* and challenged with *T. ni*. Metabolite concentrations were quantified in the harvested leaves (n = 4 independent experiments with 8 plants per treatment). Statistical significance was determined by using two-way ANOVA and multiple comparison was performed using Fisher’s LSD test. Different letters denote significant differences at *p* < 0.05. **b**, ISR against *T. ni* by *L. bicolor* is dependent on tryptophan biosynthesis via *CYP79B2/B3* (n = 3 experiments with at least 20 caterpillars per treatment). Statistical significance between treatments was determined by performing two-way ANOVA of the linear mixed model with Tukey’s test. Different letters indicate significance at *p* < 0.05 and the error bars indicate SE.

### CERK1 is necessary for *L. bicolor* induced ISR

During colonization of plant roots, beneficial *Pseudomonas* spp. both suppress^45,46^ and induce^23^ distinct subsets of defence-related genes indicating that beneficial microbes can induce and suppress PTI. To test whether *L. bicolor* induces PTI, *Arabidopsis* seedlings were treated with live and heat-killed *L. bicolor* and MAP Kinase phosphorylation was used as a readout for PTI. We found that both live and dead *L. bicolor* triggered phosphorylation of MPK3, MPK4 and MPK6 in *Arabidopsis* seedlings (Figure 4a). This observation indicates that both live and dead *L. bicolor* trigger PTI in the local tissue of *Arabidopsis*.

**Figure 4.**
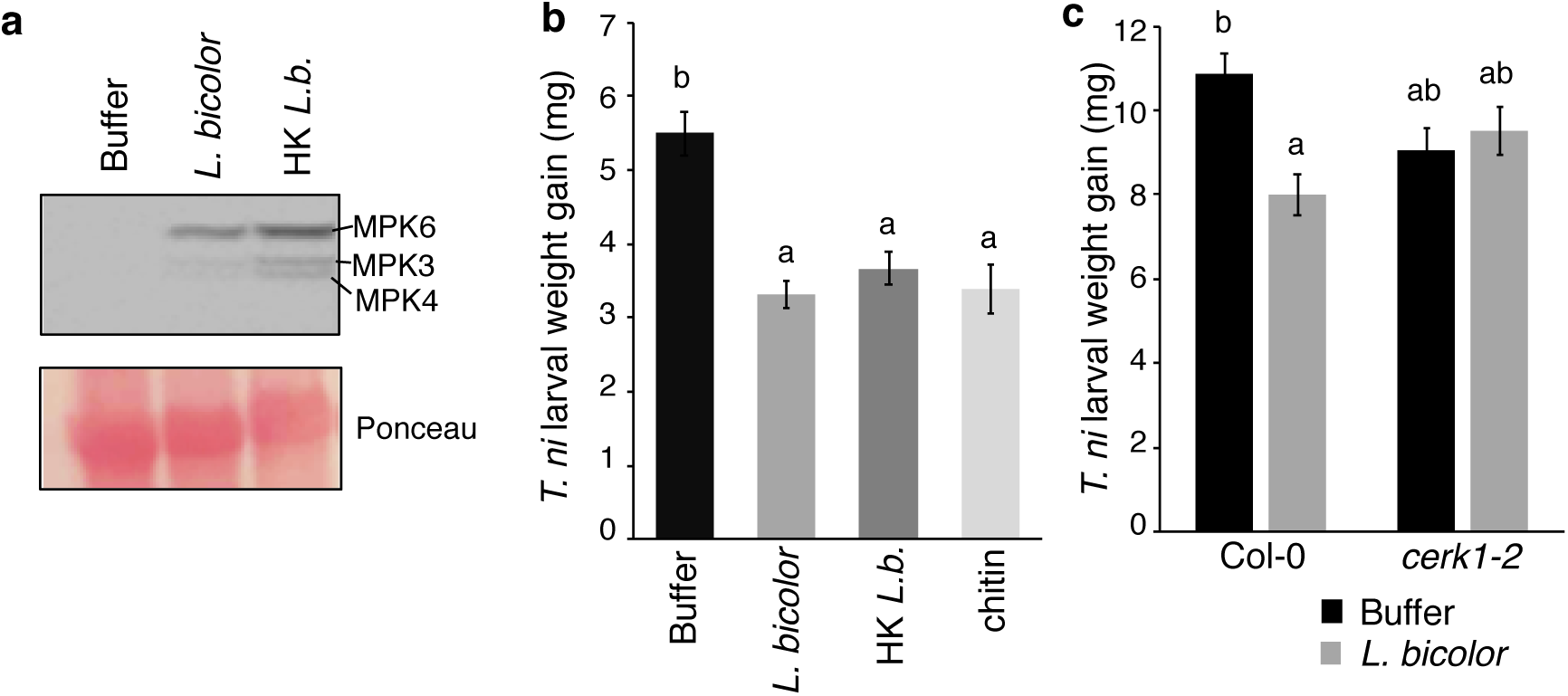
Chitin is sufficient to induce ISR and *L. bicolor* triggers ISR against *T. ni* in a *CERK1* dependent manner. **a**, Phosphorylation of MAP Kinase proteins in *Arabidopsis* seedlings treated with live and heat-killed *L. bicolor* (HK *L. b.*) for 15 mins (repeated twice with similar results). MPK6, MPK3 and MPK4 present in the samples were detected in immunoblots using α-p44/42-ERK antibody and Ponceau Red staining of the membrane. **b**, Similar to live *L. bicolor, Arabidopsis* root treatment with heat-killed *L. bicolor* and chitin resulted in significant reduction in *T. ni* larval weight gain. n = 3 experiments with at least 20 caterpillars per treatment. **c**, *L. bicolor* inoculation did not significantly affect *T. ni* growth in *cerk1-2 mutants*. n = 3 experiments with at least 20 caterpillars per treatment. Statistical significance between treatments was determined by performing two-way ANOVA of the linear mixed model with Tukey’s test. Different indicate significance at *p* < 0.05 and the error bars indicate SE.

That *L. bicolor* can induce PTI led us to ask whether *L. bicolor* triggered ISR against *T. ni* might be the result of fungal MAMP perception by plant roots. To test this hypothesis, *Arabidopsis* seedlings were inoculated with heat-killed *L. bicolor* and challenged with *T. ni*. We found that treating *Arabidopsis* roots with heat-killed *L. bicolor* caused a 33% reduction in *T. ni* larval weight gain (Figure 4b, *p* < 0.0001). This indicates that live *L. bicolor* is not necessary to trigger ISR and that MAMP perception may underlie *L. bicolor-*mediated ISR against *T. ni*.

Chitin is a structural component of the fungal cell wall and can induce PTI in plants via the CERK1 receptor^7,47^. We tested whether chitin is sufficient to trigger ISR against *T. ni* when applied to roots. We observed that chitin treatment of *Arabidopsis* roots resulted in a 38% reduction in *T. ni* larval weight gain relative to buffer-treated plants (Figure 4b, *p* < 0.0001). This indicates that chitin is sufficient to trigger ISR against *T. ni*. To test whether plant perception of chitin is necessary for ISR by *L. bicolor*, ISR experiments were performed with the chitin receptor mutant, *cerk1-2*. We found that the treatment of *cerk1-2* mutant roots with *L. bicolor* did not inhibit *T. ni* larval weight gain (Figure 4c). Thus, ISR by *L. bicolor* is dependent on *CERK1*, which indicates that ISR by *L. bicolor* is due to chitin perception by *Arabidopsis* roots.

Like *Pseudomonas*-triggered ISR against herbivores, L. *bicolor*-mediated ISR against *ni* depends on JA signalling and of SA biosynthesis (Figure 1)^17^. Many microbes that trigger ISR against herbivores can affect defence responses against other pests and pathogens. For instance, *P. simiae* WCS417 triggers ISR against bacterial pathogens while others trigger induced systemic susceptibility (ISS)^17^. To determine if *L. bicolor* can protect *Arabidopsis* from other threats, *Arabidopsis* plants were treated with *L. bicolor* and infected with *Pto*. When Col-0 leaf disks were harvested and CFUs were counted, we found that *L. bicolor* (both live and dead) and chitin treatment induced systemic susceptibility (ISS) to *Pto* (Figure 5a; *p* < 0.001). As with *L. bicolor*-induced ISR against *T. ni*, we found that *L. bicolor* triggered ISS against *Pto* is dependent on *CERK1* (Figure 5b). These data indicate that in some instances, ISR against *T. ni* and ISS against *Pto* might be due to induction of PTI in plant roots.

**Figure 5.**
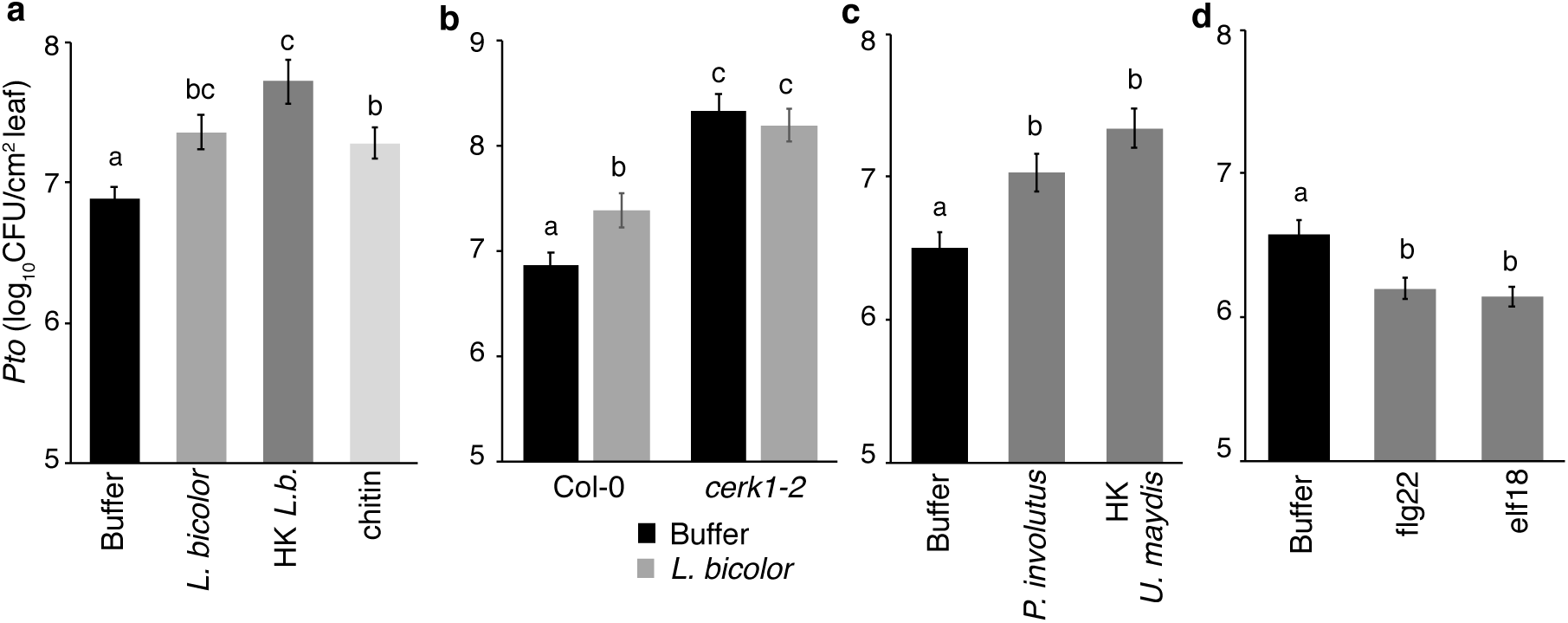
*L. bicolor* and chitin perception induce systemic susceptibility (ISS) in *Arabidopsis* against a bacterial pathogen. **a**, *Arabidopsis* plants inoculated with chitin, and live and dead *L. bicolor* had significantly increased *Pto* growth relative to mock-treated plants. **b**, *L. bicolor* inoculation did not significantly affect *Pto* infection in *cerk1-2* mutants. **c**, *P. involutus* and heat-killed (HK) *U. maydis* D132 treatment of roots induced ISS to *Pto* infection. **d**, flg22 and elf18 treatment of roots triggered ISR in *Arabidopsis* seedlings against *Pto*. Data show log_10_ (*Pto* CFU/mL) with n = 4 experiments for **a** and **d**, n = 3 experiments for **b** and **c** with 12 leaves per treatment. Statistical significance between treatments was determined by performing ANOVA. Different letters indicate significance at *p* < 0.05 and the error bars indicate SE.

If chitin is sufficient to induce resistance against *T. ni* and susceptibility against *Pto*, treatment of Col-0 roots with *Paxillus involutus*, another ECM fungus or a pathogen like *Ustilago maydis* should also result in ISS against *Pto*. However, *U. maydis* can cause necrosis and stunting of *Arabidopsis*^48^. To avoid fungal pathogenicity confounding the data interpretation, *Arabidopsis* roots were treated with heat-killed *U. maydis* D132. We found that *Arabidopsis* plants treated with live mycelium of the ECM fungus *P. involutus* and heat-killed *maydis* D132 also triggered ISS against *Pto* [Figure 5c; *p* < 0.01 (*P. involutus*) & *p* < 0.001 (*U. maydis* D132)]. To test if additional MAMPs also induce ISS, *Arabidopsis* roots were treated with flg22, elf18 and leaves were infiltrated with *Pto*. In contrast to chitin, flg22 and elf18 treatments significantly reduced *Pto* CFUs in *Arabidopsis* leaves (Figure 5d, *p* < 0.0097 and *p* < 0.00016 respectively). These data show that perception of chitin in the roots modulates systemic immunity against both insect herbivores and biotrophic pathogens and that not all MAMPs have the same effect on systemic immunity.

## DISCUSSION

The evolutionary history of some symbiotic plant-microbe interactions is well defined, while others remain more nebulous. Nitrogen-fixing rhizobia and AM and ECM fungi engage in highly specific signal exchange to initiate nutritional symbiosis with their plant hosts^30^. Whether similar co-evolution and symbiotic specificity are necessary for ISR triggered by mycorrhizal fungi and rhizobacteria is unknown. We used a non-mycorrhizal model consisting of *Arabidopsis* and the ECM fungus, *L. bicolor* to address the specificity of ISR. We found that *L. bicolor* can trigger ISR in non-mycorrhized host, indicating that ISR can be independent of mutualistic symbiosis (Figure 1a). This is consistent with ISR-induction on distinct hosts from that which a strain was originally isolated^20–23^.

We found that *L. bicolor* treatment of *Arabidopsis* roots results in a local PTI response (Figure 3a). Our data show that chitin, a fungal MAMP, is sufficient to trigger ISR against *T. ni* and ISS against *Pto* (Figure 4b). The bacterial MAMPs flg22 and lipopolysaccharide (LPS) have previously been shown to be sufficient to trigger ISR against bacterial pathogens^49^ and both flagellin and WCS417 trigger overlapping transcriptional responses in *Arabidopsis* roots^23^. Interestingly, in addition to a role for chitin perception and immunity signalling, AM symbiosis in rice is also dependent on CERK1^50^. While not all MAMPs result in the same effects on systemic defences, collectively these observations suggest that in some cases, ISR, a potentially beneficial consequence of plant-microbe interactions, might be due to MAMP exposure rather than host colonization.

Thus far, ISR has been studied in mutualistic symbioses (AM and ECM fungi), or where the evolutionary history of the interaction is unknown. Our findings show that plants also respond to non-adapted microbes resulting in ISR against herbivory. This suggests that testing isolates known to induce ISR, even if on a phylogenetically distant plant host, may be a rational approach to identifying ISR-inducing strains. These findings also suggest that chitin from a variety of sources may be useful in promoting ISR in agricultural plants against herbivores.

## MATERIALS AND METHODS

### Plant growth conditions

*Arabidopsis* seeds used in this study include the wildtype accession Columbia-0 (Col-0) and the mutants *coi1-16*^51^, *npr1-1*^52^, *sid2-2*^53^, *npr3-2/npr4-2*^54^, *cyp79b2/b3*^55^ and *cerk1-2*^7^. Seeds were sterilized with 70% ethanol for 1 min, 10% bleach for 2.5 min and washed thrice with sterile distilled water. Surface sterilized seeds were cold stratified at 4 °C for two days and planted on Jiffy-7 Horticulture peat pellets. Trays containing the pellets were covered with clear domes to maintain high humidity during seed germination. One-week-old seedlings were thinned to one seedling per pellet. The plants were watered twice a week and grown under 12 hours of light (80-100 µE) at 23 °C, 12 hours of dark at 20 °C, 70-80% relative humidity for the duration of the experiment.

### Culturing *Laccaria bicolor*

*Laccaria bicolor* (monokaryotic strain CII-H-82, S238N) was cultured on solid modified melin norkrans (MMN) medium (Composition (g/L): glucose 10.0, (NH_4_)_2_C_4_H_4_O_6_ 2.5, (NH_4_)_2_SO_4_ 0.25, KH_2_PO_4_ 0.5, MgSO_4_.7H_2_O 0.15, CaCl_2_.2H_2_O 0.05, NaCl 0.025, 0.1% thiamine-HCl 0.1 mL and 1% FeCl_3_.6H_2_0 1mL; pH 5.2 - 5.4) at 23-26 °C in the dark^56^. Three-week-old colonies were dissected using a sterile scalpel and at-least 25 agar plugs (0.25 cm^2^) were inoculated per 200 mL of liquid MMN medium. Media with agar plugs were incubated at 23 °C in the dark with shaking at 100 rpm. Mycelium that grew from the fungal plugs after 3 weeks was homogenized using a sterile mechanical shearer for less than 10 seconds. The homogenized fungal solution was poured into sterile 50 mL FALCON polypropylene conical tubes (falcon) and screw caps were partially tightened. The falcon tubes with *L. bicolor* were returned to the shaker at 23 °C in the dark for two days.

### Fungal inoculation and chitin treatment of plant roots

*L. bicolor* cultures in tubes were centrifuged at 4500 rpm for 5 min at 4 °C. The supernatant was discarded and the hyphae in the pellet was re-suspended in 10 mM magnesium sulphate (MgSO_4_) buffer. The optical density of the solution was measured at 600 nm using 10 mM MgSO_4_ as the blank. An inoculum with OD_600_ of 0.1 was used for *Arabidopsis* root treatment. 9-days old *Arabidopsis* seedling roots were inoculated with 2 ml of the *L. bicolor* solution or 10 mM MgSO_4_ for the control plants. *Paxillus involutus* culture was prepared as described above for *L. bicolor* in MMN medium and 2 mL of OD_600_ = 0.1 was inoculated on 9-days old *Arabidopsis* seedling roots.

Dead *L. bicolor* and *Ustilago maydis* D132 were prepared by heat treating a 20-fold concentration (OD_600_ of 2) of the *L. bicolor* and *U. maydis* inoculum in a water bath at 65-85 °C for 25 mins. The heat-killed cultures were inoculated on MMN and PDA agar media respectively and incubated at 23 °C in the dark to check for survival. After cooling the inoculum to room temperature, plant roots were treated with 2 mL of the heat-killed *L. bicolor* and *U. maydis*. A chitin stock (10 mg/ml) was prepared using chitin from shrimp shells (Sigma Aldrich), autoclaved for 30 mins and centrifuged to collect the supernatant^45^. Chitin treatment solution (500 µg/ml) was diluted from the stock using 10 mM MgSO_4_ and 2 ml was pipetted onto the soil surrounding each individual seedling.

### Caterpillar feeding experiments

ISR experiments were done by challenging control and *L. bicolor* treated plants with the cabbage looper, *Trichoplusia ni*^17,57^. Fresh batches of *T. ni* eggs were obtained from Natural Resources Canada with instructions that the eggs should be collected over 24 hours and shipped immediately to allow for synchronous hatching. The eggs were incubated at 23 °C in the same light regime as the plants to synchronize them with the plant circadian rhythm. First larval instars emerged 2 days after incubation. A single larva was placed on a single plant using a fine paint brush and the entire plant was covered with a breathable nylon mesh net. The hatchlings could feed on the plant for one week, after which the larvae were collected and weighed to the nearest 0.1 mg. The initial weight is negligible, and so the final weight represents the weight gain over the 1 week of feeding^58^. Each treatment or plant genotype was tested in at least 3 independent experiments (performed on different days and different batches of plants). A minimum of 20 plants per treatment per treatment were used.

For phytohormone, metabolite and qRT-PCR analyses, each plant was challenged with two *T. ni* larvae. Untreated controls were netted without *T. ni* larva and incubated under same conditions. 8 plants were used for every treatment and one leaf from each plant per treatment was harvested after 24 hours, pooled together and immediately frozen in liquid nitrogen. Samples were stored at -80 °C until processing. Only leaves with visible feeding damages were collected for *T. ni* treatment samples. Samples were collected from at least 4 independent experiments.

### *Pseudomonas syringae* pv. tomato DC3000 (*Pto*) infection assays

*Pto*^59^ infections were performed as described^60^. *Pto* inoculum was prepared by diluting an overnight culture to OD_600_ = 0.0002 using 10 mM MgSO_4_. 5-weeks old plants were watered and covered with humidity dome for a minimum of an hour. Three leaves per plant were marked using a sharpie and infiltrated with *Pto* inoculum using a blunt 1 ml syringe. Trays were covered with humidity dome and placed back under plant growth conditions. Leaf disks (0.9 cm diameter) were collected from 2 infected leaves per plant (six plants per treatment) at two days post infection. The leaf disks were homogenized, serially diluted and plated on Luria Bertani (LB) medium with rifampicin (50 µg/ml). Colony forming units (CFU) were counted 2 days after incubation at 29 °C and experiments were performed at least 3 independent times.

### *Arabidopsis* treatment with *L. bicolor* on plates

For RNAseq experiments with plants grown on solid media, 20 surface-sterilized, stratified *A. thaliana* (Col-0) seeds were germinated on MS medium agar plates^61^. After 7 days, the roots were covered with cellophane strips containing pre-cultivated mycelium of *L. bicolor*. The controls were covered with cellophane strips. After two days of exposure, the leaves which were not in contact with the fungus were harvested and frozen at -80 °C. The leaves from 5 plates were pooled. The experiment was repeated five times independently for RNA extraction and sequencing.

### RNA extraction and qRT-PCR analyses

RNA was extracted from samples collected as described above by using Qiagen RNAeasy extraction kit. The extracted RNA was quantified using Nanodrop. TURBO DNAse (Ambion) was used to remove DNA contamination from RNA samples. Single cDNA synthesis was performed using 1µg DNA-free RNA using Superscript III (Invitrogen) and Oligo dT primers in a 40 µL volume. Gene expression data was obtained by running quantitative PCR reactions with 10 µL reaction volume containing 0.5 µL cDNA, 1 µL of 5 µM primer mix containing forward and reverse primers (2.5 µM each) and 2x PowerUpTM SYBR Green master mix (Thermo Fischer). Expression values were normalized to the housekeeping gene *EIF4A*^45^ and samples from 5 biological replicates were tested. The other genes and the respective primers used in this analysis are listed in Supplementary 2.

### RNAseq and transcriptomics analyses

Quality of RNA extracted from the samples was checked using a Bioanalyzer (Agilent 2100). The RNA integrity numbers (RIN) ranged from 6.8 – 8.2 (Supplementary Tables 3 and 4). Library construction and sequencing were conducted at Chronix Biomedical (Chronix Biomedical, Inc., Göttingen, Germany). RNA libraries were prepared using the TrueSeq RNA Library Prep Kit (Illumina). Single-end reads were sequenced with a length of 75 bp on an Illumina HighSeq 2000.

Sequencing yielded 16 to 23 million reads per sample (Supplementary Tables 3 and 4). Raw sequence data was processed with the FASTX or FASTp toolkit^62^. Using FASTQ Trimmer, all nucleotides with a Phred quality score below 20 were removed from the ends of the reads, and sequences smaller than 38 bp or sequences with a Phred score below 20 for 10% of the nucleotides were discarded by the FASTQ Filter; adapter sequences and primer sequences were trimmed with the FASTQ Clipper (http://hannonlab.cshl.edu/fastx_toolkit/). Read numbers per sample after processing remained between 16 and 22 million (Supplementary Tables 3 and 4). The raw data have been deposited in https://www.ebi.ac.uk/arrayexpress/experiments/E-MTAB-8544 (gnotobiotic experiments) and https://www.ebi.ac.uk/arrayexpress/experiments/E-MTAB-8523 (soil experiments).

The processed sequences were mapped against the *Arabidopsis thaliana* transcriptome TAIR10 (downloaded from www.arabidopsis.org^63^ using Bowtie 2^64^. Bowtie mapping files were summarized to transcript count tables in R. To find transcripts with significantly increased or decreased abundance, the DEseq2 package^65^ implemented in R^66^ (R Core Team 2017) was used.

### Phytohormone and metabolite measurements

Extraction of phytohormones was carried out from the samples described above (Caterpillar feeding experiments) with methyl-tert-butyl ether (MTBE)^67^. Reversed phase separation of constituents was performed as previously described using an ACQUITY UPLC® system (Waters Corp., Milford, MA, USA) equipped with an ACQUITY UPLC® HSS T3 column (100 mm x 1 mm, 1.8 µm; Waters Corp., Milford, MA, USA). Nanoelectrospray (nanoESI) analysis was carried out as described in^67^ and phytohormones were ionized in a negative mode and determined in a scheduled multiple reaction monitoring mode with an AB Sciex 4000 QTRAP® tandem mass spectrometer (AB Sciex, Framingham, MA, USA). Mass transitions are described in Supplementary Table 5.

### Mitogen-activated protein kinase (MAPK) experiments

Two sterile seeds were germinated per well in a 24 well plate with liquid Murashige & Skoog (MS) medium and 0.5% sucrose. The media was replaced with fresh MS + sucrose medium on day 7 and with water on day 14. Elicitor and treatment solutions were prepared with sterile distilled water to the final concentration of *L. bicolor* (OD_600_ = 0.2) and 20x-dead *L. bicolor*. 15-day-old seedlings were treated with 500 µl of *L. bicolor* treatments. Seedlings were treated with for 15 minutes and immediately frozen in liquid nitrogen after 15 minutes of treatment. The samples were stored at -80 °C before being weighed. Samples were homogenized while frozen prior to protein extraction. MAP Kinase activation in the samples were analyzed according to^68^. Separated proteins were quantified by staining with Ponceau Red and de-stained using stripping buffer.

### Statistical analyses

Statistics were computed in R^66^. For the caterpillar feeding experiments, linear mixed-effect models were applied to log-transformed *T. ni* larval weight gain from all experiment repetitions. For the *Pto* CFU/mL experiments, linear mixed-effect models were applied to the untransformed data from all experiment repetitions. Experiment repetition was applied as random effect (function ‘lmer’ from package ‘lme4’^69^). Normal distribution and homogeneity of variance was verified by visual inspection of the residual plots. Statistical significance was determined by performing two-way ANOVAs on the linear mixed-effects model fit to the data (function ‘Anova’ from package ‘car’,^70^. Homogeneous subsets were determined by a post-hoc test (Tukey’s HSD, function ‘glht’ from package ‘multcomp’^71^).

For the phytohormone, metabolome and qRT-PCR analyses, the data obtained from the treatments were normalized to mock treated samples. Linear models were applied to the data. Normal distribution and homogeneity of variance was verified by visual inspection of the residual plots. Statistical significance was determined by performing two-way ANOVAs on the linear models. Homogeneous subsets were determined by a post-hoc test, Fisher’s LSD for phytohormone and metabolite analyses, and Tukey’s HSD for qRT-PCR data. RNAseq analyses were performed using the ‘DESeq2’ package^65^.

## ACKNOWLEGDEMENTS

This work was supported by funding in the frame of the International Research Training Group 2172: PRoTECT – Plant Responses To Eliminate Critical Threats by the Deutsche Forschungsgemeinschaft (DFG) hosted by the University of Goettingen (Germany), for the Service Unit for Metabolomics and Lipidomics (DFG, INST 186/822-1) to I. F. and the University of British Columbia (Canada). This work was also supported by a Natural Sciences and Engineering Research Council (NSERC) Discovery Grant (NSERC-RGPIN-2016-04121) awarded to C.H.H. Y.L was supported by an NSERC CREATE-PRoTECT award and a Chinese Graduate Scholarship Council Award. We thank K. Ziesing and Sabine Freitag for technical assistance and Volker Lipka and Sina Barghahn for support with MAPK assays and for critical reading of the manuscript.

