## Supplementary Figures and Tables for "Ectomycorrhizal fungi induce systemic resistance against insects on a non-mycorrhizal plant in a CERK1-dependent manner"

**a**

| Genotypes in experiment | Figures | Location | Date | Average Larval Weight (mg) |
|---|---|---|---|---|
| *coi1-16, npr1-1, sid2-2* | Figure 1a | UBC | 2017-08-26 | 5.34 |
| | | UBC | 2017-11-15 | 6.94 |
| | | UBC | 2017-12-06 | 4.38 |
| | | UBC | 2018-01-17 | 4.17 |
| *npr3-2/4-2, cerk1-2* | Figure 1b and 4c | UBC | 2018-03-17 | 10.23 |
| | | UBC | 2018-03-17 | 12.13 |
| | | UBC | 2018-03-23 | 10.30 |
| chitin, HK *L.b.* | Figure 4b | Goettingen | 2018-08-21 | 2.93 |
| | | Goettingen | 2018-09-25 | 9.36 |
| | | Goettingen | 2018-10-05 | 2.99 |
| *cyp79b2/b3* | Figure 3b | Goettingen | 2018-11-19 | 3.07 |
| | | UBC | 2019-03-02 | 2.56 |
| | | Goettingen | 2019-05-23 | 1.72 |

**b**

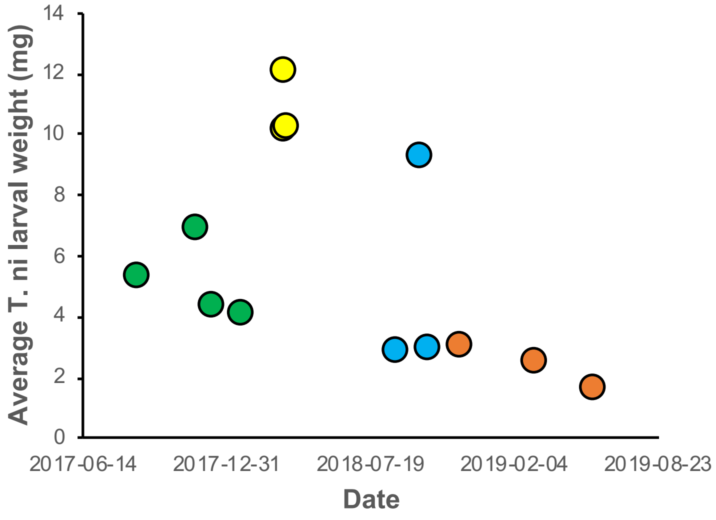


**Supplementary Figure 1. Variation in control *T. ni* larval weight over time and by location.** We noted a significant replicate effect in the weight of *T. ni* larvae feeding on buffer-treated *Arabidopsis* Col-0 control plants. **a,** average weights of the controls by date and location. **b,** the average control larval weight over time and by experiment.

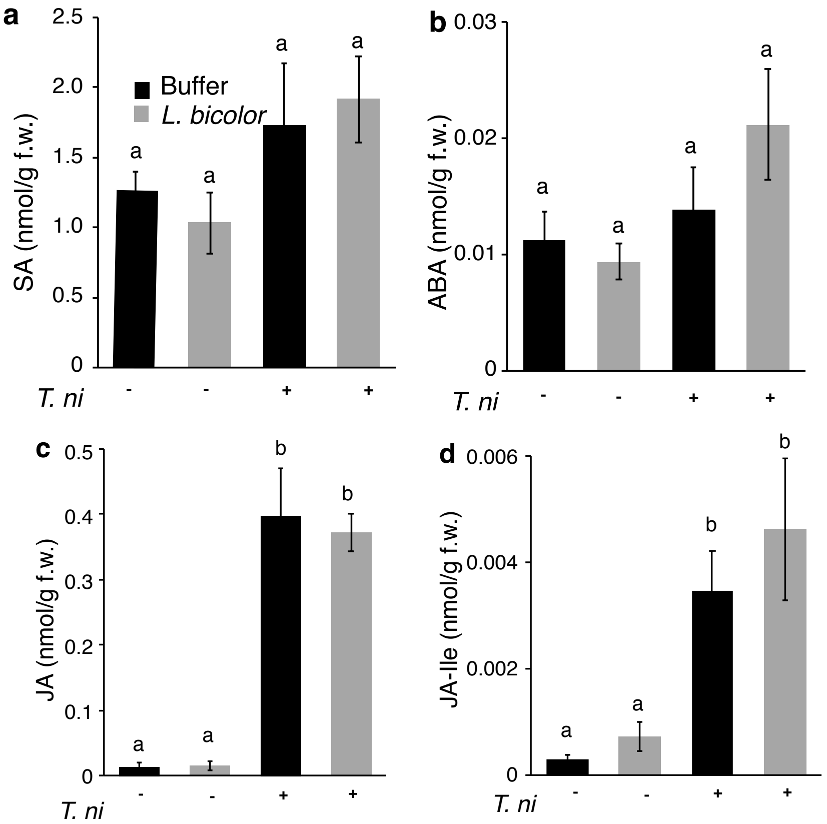


**Supplementary Figure 2. *L. bicolor* treatment of roots does not significantly increase the levels of phytohormones in *Arabidopsis* leaves.** Leaves were harvested from *Arabidopsis* Col-0 plants treated with buffer or *L. bicolor* and challenged with *T. ni*. The concentrations of **a,** SA, **b,** ABA **c**, JA and **d,** JA-Ile d were measured using GC-MS. Mean concentrations of the phytohormones in samples collected from six independent experiments (n = 6 ± SE) are shown. Statistical significance was determined using two-way ANOVA and Fisher's LSD test. Different letters denote significant differences at $p < 0.05$.

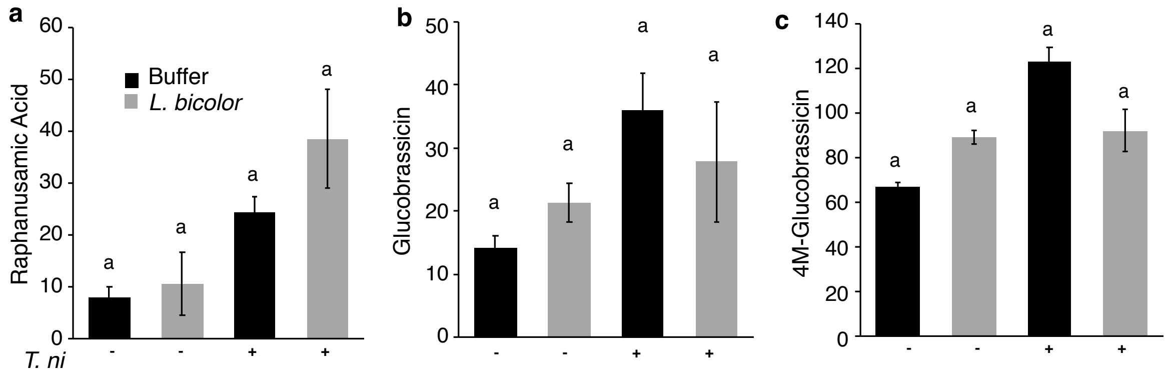


**Supplementary Figure 3. The effect of *L. bicolor* and *T. ni* on the accumulation of secondary metabolites in *Arabidopsis* leaves.** Leaves were harvested from *Arabidopsis* Col-0 plants treated with buffer or *L. bicolor* and challenged with *T. ni*. The concentrations of **a**, raphanusamic acid **b,** Glucobrassicin **c,** and 4M-Glucobrassicin were measured using GC-MS. Arbitrary units are shown. Mean concentrations as a proportion of the total metabolic of the metabolites (in samples collected from 6 independent replicates (n = 6 ) ± SE are shown. Statistical significance was determined by using two-way ANOVA and Fisher's LSD test with $p < 0.05$.

### Supplementary Tables

| Candidate gene | Functionality | Forward and reverse primers | Reference |
|---|---|---|---|
| *EIF4A* | House-keeping gene | 5′-GCAGTCTCTTCGTGCTGACA-3′ and 5′ TGTCATAGATCTGGTCCTTGAA-3′ | 1 |
| *PR1* | SA pathway | 5′-ACACCTCACTTTGGCACATC-3′ and 5′-GAGTGTGGAAAACGCAAAGA-3′ | 1 |
| *PR2* | SA pathway | 5′-CCTTCTCGGTGATCCATTCT-3′ and 5′-AGTGTGGAAAACGCAAAGACT-3′ | 1 |
| *GST6* | SA pathway | 5′-CCATCTTCAAAGGCTGGAAC-3′ and 5′-TCGAGCTCAAAGATGGTGAA-3’ | 1 |
| *MYC2* | JA pathway | 5’-AGATAAAACCGCCGGAGAAT-3’ and 5’-TACCGTTTGCTGGCTTTCTT-3’ | 2 |
| *VSP1* | JA pathway | 5′-CTCAAGCCAAACGGATCG-3’and 5′-TTCCCAACGATGTTGTACCC-3’ | 1 |
| *VSP2* | JA pathway | 5’-TCAGTGACCGTTGGAAGTTGTG-3’ and 5’-GTTCGAACCATTAGGCTTCAATATG-3’ | 3 |
| *ERF1* | JA/Et pathway | 5’-ATTCTTTCTCATCCTCTTCTTCT-3’ and 5’-CGAATCTCTTATCTCCGCCG-3’ | 4 |
| *ORA59* | JA/Et pathway | 5’– AAGGGATAAGAGTGTGGCTTGGGA-3’ and 5’- CTTTCAAAGCGAAAGCCGCCTGAT-3’ | 5 |
| *PR4* | JA/Et pathway | 5’-GAGAATAGTGGACCAATGCAG-3’ and 5’-GTAGACCGATCGATATTGACCT-3’ | 6 |
| *PDF1.2* | JA/Et pathway | 5’-AATGAGCTCTCATGGCTAAGTTTGCTTCC-3’ and 5’-AATCCATGGAATACACACGATTTAGCACC-3’ | 7 |

**Supplementary Table 2.** List of candidate genes and the primer sequences used for qRT-PCR analysis

| Sample ID | Treatment | Sample number | RIN Value | Raw | Trimmed/Filtered | Mapped | % Mapped |
|---|---|---|---|---|---|---|---|
| RCE3_K | Col-0 (1) | S1 | 7.7 | 22,120,615 | 21,823,151 | 21,067,585 | 96.54 |
| RCE3_Lb | Col-0 + *L. bicolor* (2) | S2 | 4.9 | 21,576,771 | 21,282,687 | 20,630,221 | 96.93 |
| RCE4_K | Col-0 (1) | S3 | 8 | 19,916,587 | 19,660,941 | 19,056,023 | 96.92 |
| RCE4_Lb | Col-0 + *L. bicolor* (2) | S4 | 7.9 | 20,633,797 | 20,367,512 | 19,743,724 | 96.94 |
| RCE5_K | Col-0 (1) | S5 | 6.8 | 20,501,270 | 20,221,234 | 19,604,781 | 96.95 |
| RCE5_Lb | Col-0 + *L. bicolor* (2) | S6 | 6.9 | 20,034,091 | 19,771,853 | 19,129,605 | 96.75 |
| RCE6_K | Col-0 (1) | S7 | 7.1 | 21,459,527 | 21,179,595 | 20,521,235 | 96.89 |
| RCE6_Lb | Col-0 + *L. bicolor* (2) | S8 | 7 | 20,336,169 | 20,059,707 | 19,404,898 | 96.74 |
| RCE7_K | Col-0 (1) | S9 | 6.9 | 20,090,234 | 19,830,895 | 19,218,468 | 96.91 |
| RCE7_Lb | Col-0 + *L. bicolor* (2) | S10 | 6.8 | 20,421,182 | 20,169,941 | 19,529,256 | 96.82 |

**Supplementary Table 3.** Raw read numbers and mapped reads per sample for RNAseq experiment with gnotobiotic plants (shown in Figure 2a and 2b).

| Sample ID | Treatment | Sample number | RIN Value | Raw | Trimmed | Filtered | Mapped | % Mapped |
|---|---|---|---|---|---|---|---|---|
| 10A | Col-0 + Buffer | S20 | 7.50 | 18,323,659 | 18,318,236 | 18,087,530 | 17,175,563 | 94.96 |
| 10B | Col-0 + Buffer + *T. ni* | S19 | 6.80 | 20,203,128 | 20,194,911 | 19,905,113 | 18,891,328 | 94.91 |
| 13A | Col-0 + Buffer | S18 | 8.30 | 18,217,338 | 18,211,936 | 17,967,732 | 17,326,589 | 96.43 |
| 13B | Col-0 + Buffer + *T. ni* | S17 | 7.60 | 19,716,705 | 19,707,668 | 19,428,965 | 18,724,741 | 96.38 |
| 15A | Col-0 + Buffer | S16 | 7.50 | 19,706,504 | 19,697,626 | 19,414,133 | 18,621,357 | 95.92 |
| 15B | Col-0 + Buffer + *T. ni* | S15 | 7.20 | 17,364,845 | 17,357,509 | 17,100,108 | 16,447,675 | 96.18 |
| 17A | Col-0 + Buffer | S14 | 7.70 | 16,917,427 | 16,910,467 | 16,680,497 | 15,957,944 | 95.67 |
| 17B | Col-0 + Buffer + *T. ni* | S13 | 7.70 | 17,223,761 | 17,217,803 | 16,970,427 | 16,345,331 | 96.32 |
| 18A | Col-0 + Buffer | S12 | 7.90 | 19,157,737 | 19,149,695 | 18,878,821 | 18,195,574 | 96.38 |
| 18B | Col-0 + Buffer + *T. ni* | S11 | 7.60 | 20,488,629 | 20,480,817 | 20,206,424 | 19,341,346 | 95.72 |
| 20A | Col-0 + *L. bicolor* | S10 | 7.10 | 19,809,126 | 19,801,344 | 19,532,487 | 18,648,387 | 95.47 |
| 20B | Col-0 + *L. bicolor* + *T. ni* | S9 | 7.30 | 20,883,945 | 20,875,671 | 20,587,068 | 19,616,510 | 95.29 |
| 23A | Col-0 + *L. bicolor* | S8 | 8.20 | 17,267,981 | 17,262,882 | 17,029,735 | 16,397,067 | 96.28 |
| 23B | Col-0 + *L. bicolor* + *T. ni* | S7 | 7.00 | 18,245,284 | 18,236,417 | 17,966,636 | 17,202,430 | 95.75 |
| 25A | Col-0 + *L. bicolor* | S6 | 8.10 | 17,953,224 | 17,945,646 | 17,697,924 | 17,001,164 | 96.06 |
| 25B | Col-0 + *L. bicolor* + *T. ni* | S5 | 7.20 | 20,835,382 | 20,824,065 | 20,517,536 | 19,653,407 | 95.79 |
| 27A | Col-0 + *L. bicolor* | S4 | 7.90 | 19,920,333 | 19,910,035 | 19,606,826 | 18,731,911 | 95.54 |
| 27B | Col-0 + *L. bicolor* + *T. ni* | S3 | 7.60 | 18,454,061 | 18,447,710 | 18,197,177 | 17,531,189 | 96.34 |
| 28A | Col-0 + *L. bicolor* | S2 | 7.70 | 16,605,947 | 16,599,168 | 16,363,095 | 15,671,379 | 95.77 |
| 28B | Col-0 + *L. bicolor* + *T. ni* | S1 | 7.80 | 17,832,755 | 17,824,806 | 17,580,314 | 16,905,795 | 96.16 |

**Supplementary Table 4.** Raw read numbers and mapped reads per sample for RNAseq experiment with soil growth plants (shown in Figure 2c).

| MRM Transitions | | Analyte | DP [declustering potential] | EP [entrance potential] | CE [collision energy] |
|---|---|---|---|---|---|
| Q1 | Q3 | | | | |
| 137 | 93 | SA | -25 | -6 | -20 |
| 162 | 58 | RA | -15 | -6 | -14 |
| 201 | 59 | Camalexin | -51 | -8 | -45 |
| 209 | 59 | JA | -30 | -4.5 | -24 |
| 322 | 130 | JA-Ile | -45 | -5 | -28 |
| 263 | 153 | ABA | -35 | -4 | -14 |
| 447 | 97 | Glucobrassicin | -45 | -7 | -40 |
| 477 | 97 | 4-M-glucobrassicin | -55 | -5 | -38 |

**Supplementary Table 5.** Mass transitions and corresponding conditions for identification of phytohormones and secondary metabolites shown in Figures 3a, S2 and S3.
